# Wavelength induced cultivar specific enrichment of essential amino acids and phenolics in *Amaranthus tricolor*

**DOI:** 10.64898/2026.03.28.714947

**Authors:** Shweta S. Pawar, Neeraj Joshi, Yogesh Pant, Maneesh Lingwan, Shyam Kumar Masakapalli

**Affiliations:** School of Biosciences and Bioengineering, Indian Institute of Technology Mandi, 175075, Himachal Pradesh, India; Donald Danforth Plant Science Center, St. Louis, MO, USA

**Keywords:** *Amaranthus tricolor*, metabolite profiling, metabolite enrichment, GC-MS, light wavelength, essential amino acids, phenolics

## Abstract

Light wavelengths modulate plant growth, metabolism, and physiology. *Amaranthus*, a C4 underutilized climate resilient crop with promising nutritional properties remained unexplored in terms of metabolite enrichment under monochromatic light wavelengths of visible spectrum. In current study, two cultivars of *Amaranthus tricolor* (green and red) were exposed to seven light regimes of photosynthetically active radiation (PAR; 400-700 nm): deep blue, blue, green, amber, red, deep red, far red, and their metabolic responses were captured using Gas Chromatography-Mass Spectrometry. The metabolic analysis revealed wavelength-specific reprogramming in the levels of organic acids, sugars, amino acids, fatty acids as well as phenolics. In both the green and red *Amaranthus*, branched-chain amino acids and phenylalanine, which are nutritionally essential, were significantly elevated under far-red light. While the phenolics such as caffeic acid and ferulic acid were elevated under green and deep blue light respectively in green *Amaranthus*, amber light wavelengths enhanced these phenolics in red *Amaranthus*. The study highlighted cultivar-specific metabolic rewiring triggered by specific wavelengths. Altogether, these findings provides insights into metabolic adaptation and demonstrate the ability of light wavelength to specifically enrich the targeted metabolite of nutritional relevance in *Amaranthus*. It offers strategies to improve the nutritional value of crops in controlled agriculture systems.

**Graphical Abstract:** 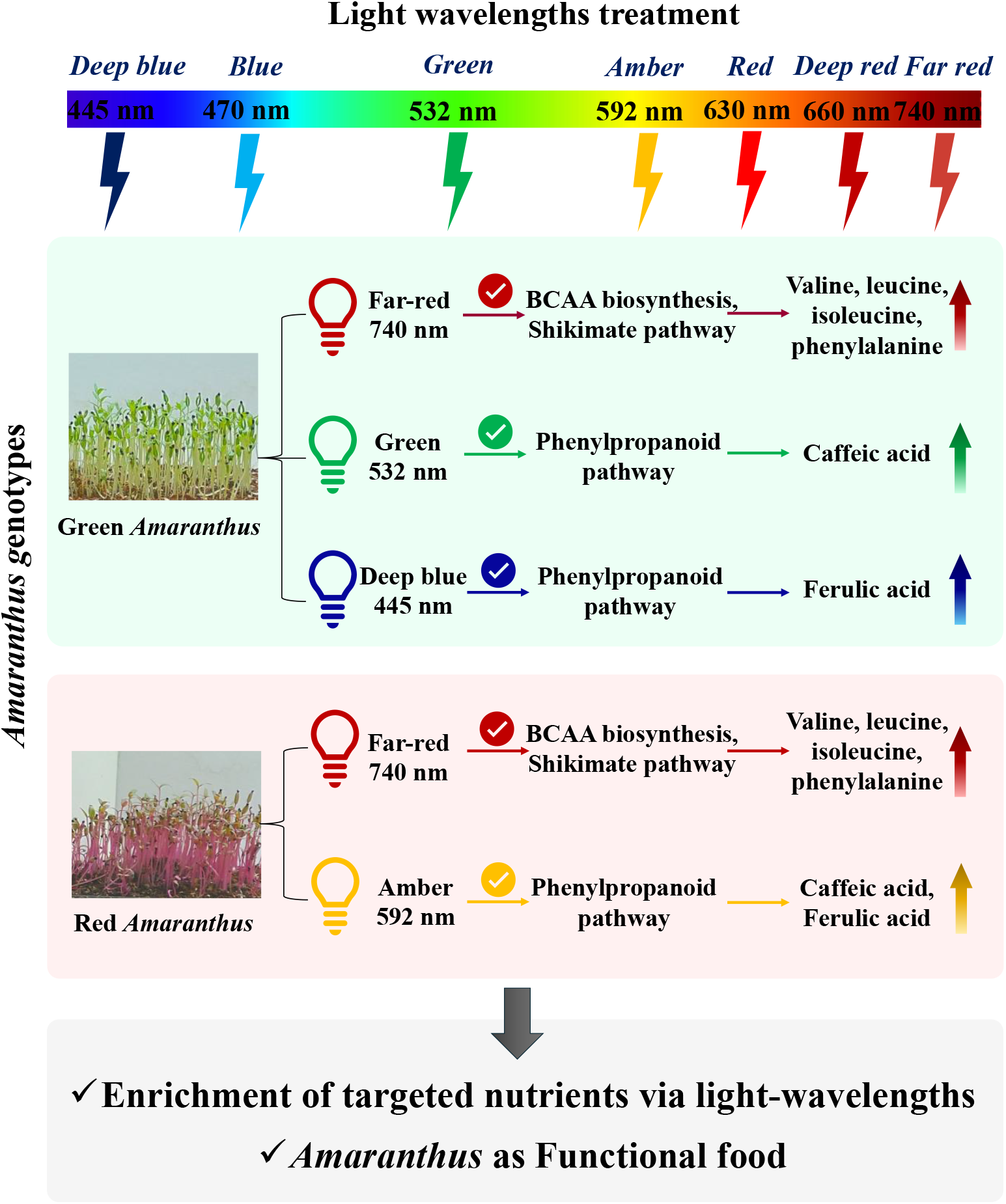

## Introduction

Light supply energy for photosynthesis and also regulates plant growth, metabolism, and physiology. Plant perceive different wavelengths of light ranging from ultraviolet to far-red through specialized photoreceptors, which play key role in regulating development and metabolism (Yadav et al., 2020). Among them, maximum photosynthesis in plants has been reported under visible light spectrum, which contains wavelengths ranging from 400 to 700 nm. However, different light wavelengths and intensities alters the metabolism of the plants leading to variation in the levels of metabolites (Thakur et al., 2025).

In indoor cultivation systems, particularly in green house, light functions as a fundamental environmental signal, supplying energy for plant development and facilitating acclimation (Song et al., 2020; Viršilė et al., 2020). Sunlight offers the full spectrum, while artificial lights provide a range of specific wavelengths that would trigger specific signalling responses in plants. The recent advancement of light-emitting diodes (LEDs) offers a promising illumination source for indoor growth chambers, as they emit a broad spectrum of light (Yeh and Chung, 2009). Light modulation at specific monochromatic light wavelengths has been shown to reprogram plant metabolism, leading to targeted enrichment of metabolites. In recent studies, significant accumulation of specialized metabolites was reported in plants under different LEDs (Alrifai et al., 2019; Mao et al., 2021; Chini Zittelli et al., 2022; Wu et al., 2024; Sae-Foo et al., 2025; Van Brenk et al., 2025).

The increasing population, changing climate and competition in the utilization of land for food and natural compounds reinforce the need for artificial growing systems (Darko et al., 2014). *Amaranthus* is an ancient C4 crop that has gained global interest due to its climate resilient traits, nutritional benefits of its grains and leafy microgreens (Sarker and Oba, 2019). While Amaranth grains are the most consumed component of the plant, its leaves serve as a rich yet underutilized resource, especially in areas where traditional green vegetables are essential for food and nutritional security (Achigan-Dako et al., 2014; Baginsky-Guerrero et al., 2026). Amaranth leaves are resilient to extreme climate conditions and offer vital nutrients like fibre, vitamins, minerals, amino acids and different bioactive substances (Sarker et al., 2020). Their composition, however, is influenced by environmental factors including light wavelength, intensity, temperature and water availability. Several leafy greens are reported as functional foods rich in amino acids like leucine, isoleucine, valine and phenolics like sinapinic acid, highlighting them as an exceptional nutrient source for addressing hidden hunger (Pant et al., 2023). The responses of green and red amaranth microgreens under different monochromatic light wavelengths of the complete visible spectrum remain unexplored. Here in this study, we have explored the wavelength and cultivar-specific metabolic responses of *Amaranthus* microgreens.

This study evaluates the impact of different light wavelengths on metabolism in *Amaranthus tricolor*. Further, investigations into cultivar-specific metabolic responses in green and red *Amaranthus tricolor* under different light wavelengths are carried out. Finally, the study focuses on identifying specific wavelengths of light that trigger the targeted enrichment of nutritional and bioactive compounds. The study established that different light conditions affect the metabolite profiles in *Amaranthus*. Further wavelength induced enrichment of specific metabolites of nutritional relevance is observed. This study lays the foundation for using *Amaranthus* microgreens as a scalable functional food, focused on enriching nutrients that offer health benefits and have industrial uses.

## Material and methodology

### Plant material and growth conditions for *Amaranthus* seedlings

The seeds of two varieties of *Amaranthus tricolor*, Pusa Kiran (green) and Pusa Lal Chaulai (red), were acquired from the Division of Vegetable Science, Indian Agricultural Research Institute, New Delhi, India. 500 mg per plate of seeds were sown in square petri plates with substrate coco peat: vermiculite (1:1) with adequate moisture content. Next, seeds were subjected to germination in dark conditions for four days maintaining 22 ± 2°C temperature as well as 60 ± 5% relative humidity (RH). They were placed in the dark for four days. After 4 days of germination in the dark, on the 5^th^ day, the seedlings were shifted to light conditions with an 8:16 h light: dark cycle up to 11^th^ day, since *Amaranthus* is a short-day plant. On the 12^th^ day, the two-leaf-stage seedlings were shifted to different wavelengths of light, including deep blue (445 nm), blue (470 nm), green (532 nm), amber (592 nm), red (630 nm), deep red (660 nm), far red (740 nm), and white light as a control (PAR region; 400-700 nm). The wavelengths of the solid-state lighting (SSL) lamps used in the study have a span of ±20 nm. The seedlings were exposed to 8 hrs of different wavelengths of light followed by 16 hrs of dark phase for the next three consecutive days (**Supplementary figure 1**). Fully grown plants were harvested on the 15^th^ day, with quenching by liquid nitrogen, as well as stored at -80°C for further processing. The samples were crushed in liquid nitrogen, then freeze-dried in a lyophilizer. Lyophilized samples were used for further analysis.

### GC-MS based metabolite profiling of green and red *Amaranthus* seedlings

The metabolite extraction of green and red *Amaranthus* seedlings was performed using 20 mg lyophilized samples in an extraction buffer containing methanol:choloroform:water in 3:1:1 ratio. 0.2 mg/ml of ribitol stock was prepared in water and 60 μl of ribitol was added per sample. Mixture was then incubated in thermoshaker for 5 minutes at 70°C at 950 rpm and then centrifuged for 10 min at RT (room temperature) at 13000 g (Lisec et al., 2006). Next, 50 μl supernatant was taken and put into another Eppendorf tube. It was dried in speedvac for further derivatization step. To each dried sample 35 μl of MeOX-pyridine was added from 20 mg/ml of stock in pyridine. After that, it was incubated for two hours at 37°C and 900 g. Each tube was then added with 49 μl of MSTFA (N-methyl-N-(trimethylsilyl)-trifluoroacetamide) and again incubated at 37°C for 10 minutes at 900 g. Later, 60 μl supernatant was transferred 0.2 ml volume new inserts (Lingwan and Masakapalli, 2022). An Agilant Technologies GC ALS-MS 5977B GC-MS, with 30 m x 250 μm x 0.25 μm HP-5MS column (5% phenyl methyl siloxane) was used for data acquisition. 1 μl of sample was injected in splitless mode at 70 eV electron ionisation. Additional program parameters, such as temperature levels and scan mass-range specifications, were established as outlined by Lingwan and Masakapalli (2022). The raw GC-MS mass spectra files were baseline corrected using MetAlign software. Additionally, by examining retention time as well as fragmentation patterns, corresponding metabolite peaks were detected with Agilent ChemStation software utilizing the NIST 17 library, with an identification score of >70% and in-house standards available.

### Metabolomics and Statistical Analysis

The relative abundances of detected metabolites were normalized using area of ribitol as an internal standard to get the relative proportions. An online tool MetaboAnalyst 6.0 was utilized to perform multivariate statistical analysis using detected compounds with their normalized abundances. Further data pre-processing was performed using specific parameters such as normalization by median, log transformation (base 10), as well as pareto scaling and PCA (Principal component analysis) plots were created (Chong et al., 2019). Pathway enrichment analysis was performed using KEGG and metaboanalyst 6.0. Normalized relative abundances of identified peaks were used to calculate fold changes of metabolites among the treatments. To determine the statistical significance or non-significance, one-way ANOVA along with Dunnett’s multiple comparison test was used with n=4 where, mean± standard deviation (SD) represents the error bars.

## Result and discussion

### Green *Amaranthus* seedlings under different light wavelengths showed distinct metabolomes

Metabolic profiling captured 52 features from soluble extracts of 14-day-old *Amaranthus* seedlings exposed to different light wavelengths (**Figure 1a**). Further, variations in metabolic profiles of green *Amaranthus* under different light wavelength were captured through multivariate statistical analysis. PCA showed different clusters of light treatments along 1^st^ two principal components (PCs), with PC1 and PC2 accounting for 30.8% as well as 19% of variation in the metabolic profiles respectively **(Figure 1b)**. PAR light treatment formed a distinct cluster, whereas far-red light treatment clustered farther away from amber, blue, deep blue and deep red regions, reflecting wavelength mediated metabolic responses. Major metabolites contributing to separation includes phenylalanine, valine, leucine, isoleucine, caffeic acid, ferulic acid, citric acid, malic acid, aconitic acid, oxalic acid etc. Previous studies highlighted short and long wavelengths activate non-overlapping metabolic networks influenced by separate photoreceptor families (phototropins, cryptochromes, phytochromes) (Chen et al., 2004; Q. Li and Kubota, 2009). This indicates variations in the metabolites could result due to the exposure to different wavelengths. Enrichment analysis highlighted, glyoxylate and dicarboxylate metabolism as the most responsive pathways followed by amino acid metabolism, TCA cycle and phenylpropanoid pathway **(Figure 1c)**.

**Figure 1.**
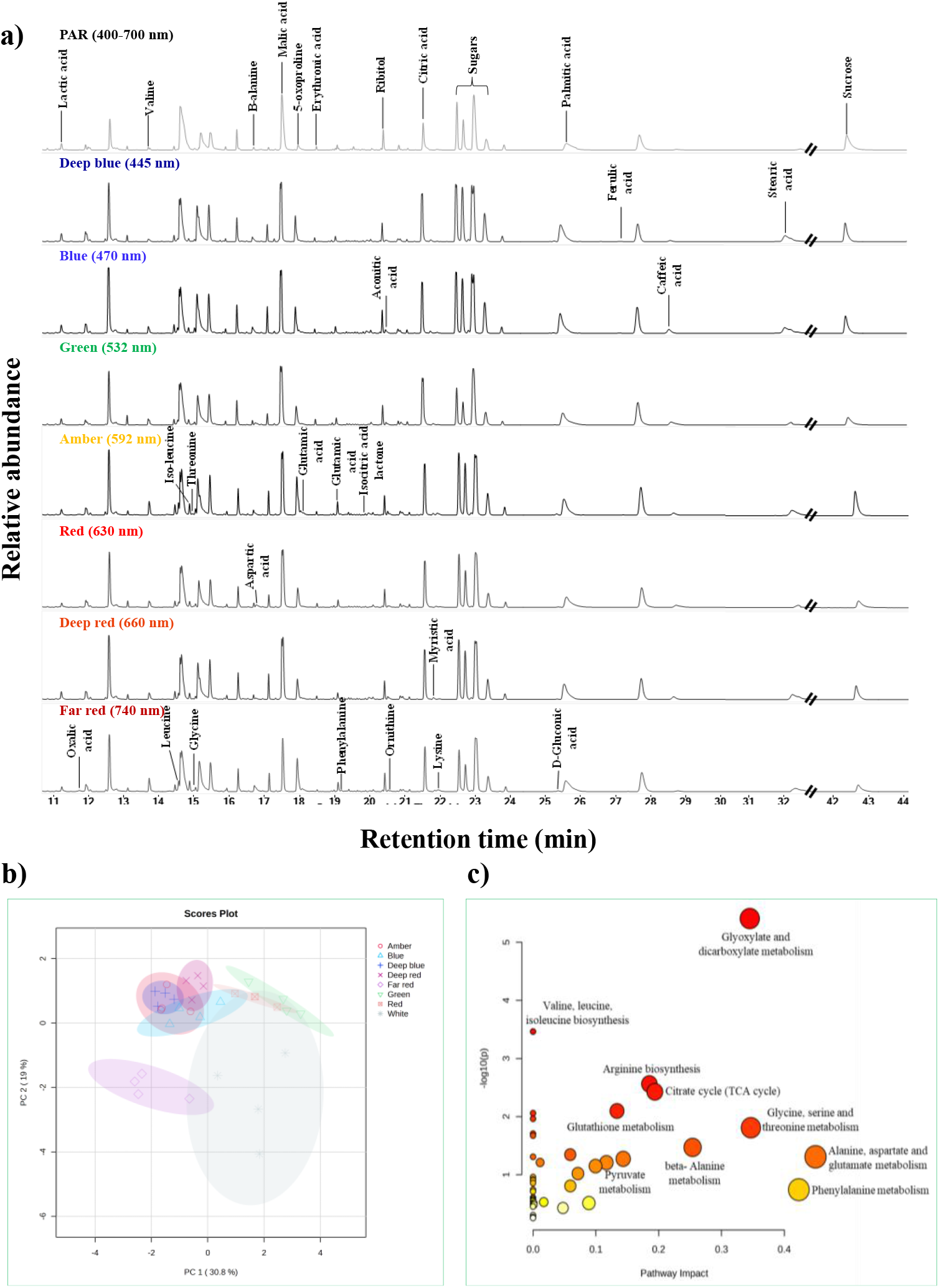
Wavelength induced metabolic responses in green *Amaranthus*. a) Overlayed GC-MS chromatograms show metabolite profiles of green *Amaranthus* subjected to PAR (White) as control and monochromatic light treatments: deep blue, blue, green, amber, red, deep red, as well as far-red. Peaks corresponding to key metabolites-including organic acids like aconitic acid, malic acid, citric acid, amino acids like lysine, threonine, leucine, isoleucine, valine, phenylalanine, fatty acids like palmitic acid, stearic acid, phenolics like ferulic acid, caffeic acid and sugars like sucrose are annotated. Changes in peak intensities illustrate wavelength-dependent shifts in metabolic abundance. b) PCA score plots of green *Amaranthus* demonstrate clear and distinct clustering under different light wavelengths. c) Pathway enrichment analysis of metabolic profiles of the significant metabolic pathways using KEGG with an impact ≥ 0.1 for green *Amaranthus* seedlings under different light treatments. In a figure, each bubble depicts a metabolic pathway; the x-axis shows the pathway impact and y-axis shows the pathway enrichment. The key pathways impact and enrichment values are shown by the color and size, respectively.

### Different light wavelength treatments also altered the metabolite profiles of red *Amaranthus* seedlings

Further to investigate whether there are any cultivars or pigment specific responses, red *Amaranthus* was subjected to different light wavelengths. The metabolite profiling highlighted wavelength dependent modulation of metabolism also in the red *Amaranthus* seedlings **(Figure 2a)**. The levels of these metabolites fluctuated under different light wavelengths conditions and revealed distinct metabolic responses compared to the green *Amaranthus* seedlings **(Figure 1)**. The metabolomics of red *Amaranthus* showed different clusters of light treatments represented by the PCA. The PC1 as well as PC2 explained 34.4% as well as 21.9% of variation in metabolic profiles respectively **(Figure 2b)**. Far red and deep blue light wavelengths clustered farther away from other treatments indicating strong wavelength dependent metabolic differences. As the number of metabolites are same in green and red Amaranthus seedlings, KEGG enrichment analysis showed similar responses as that of green *Amaranthus* with major metabolite groups associated with glyoxylate and dicarboxylate cycle, amino acid metabolism, TCA metabolism as well as phenylpropanoid metabolism under different light treatments **(Figure 2c)**. Red *Amaranthus* have more specialized metabolism compared to green *Amaranthus*, with high response of phenylpropanoid metabolism to different light wavelengths. The metabolic observations are further explained below to get insights into the effect of different light wavelengths on green and red cultivars.

**Figure 2.**
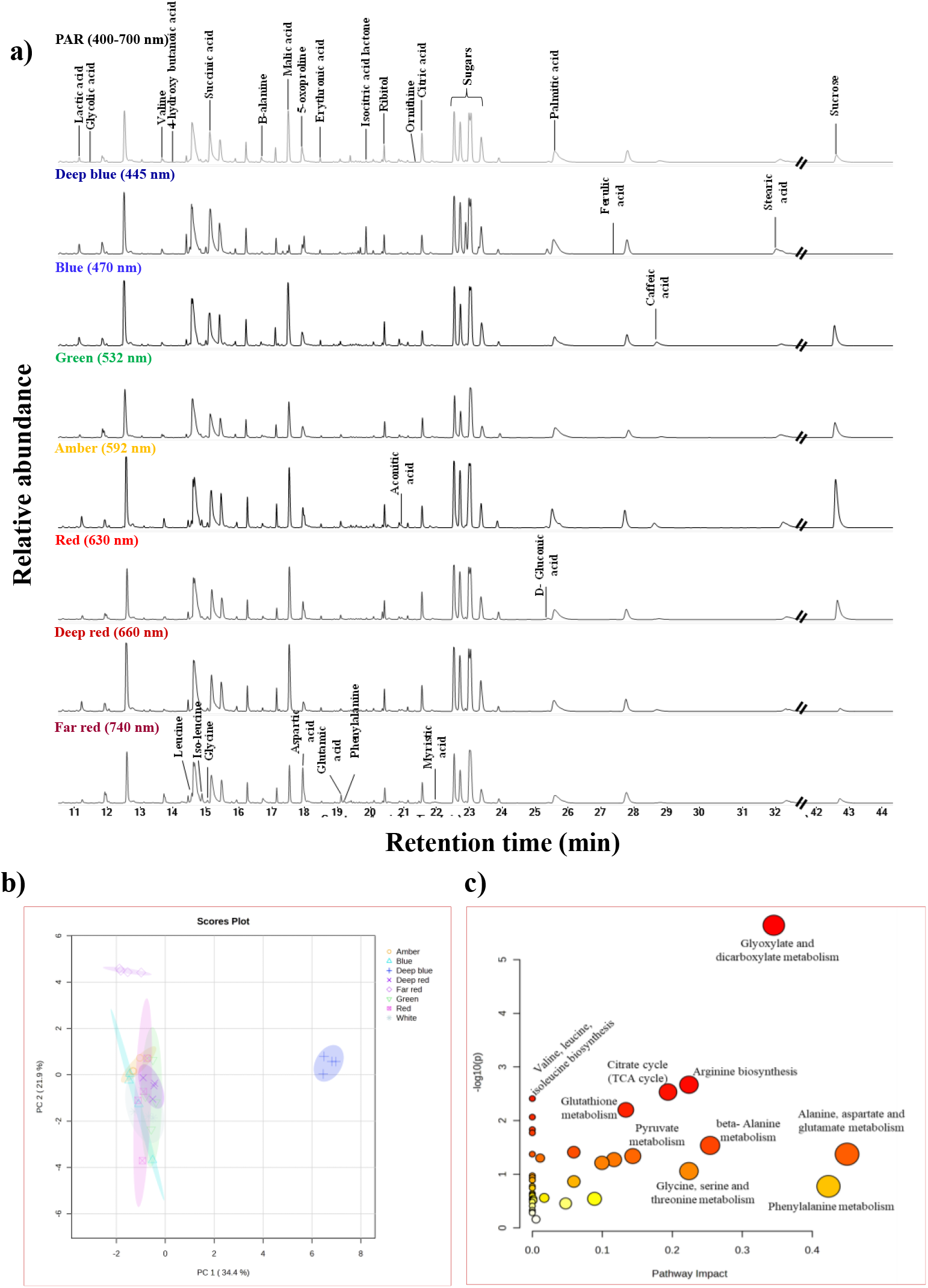
Wavelength induced metabolic responses in red *Amaranthus*. a) Overlayed GC-MS chromatograms show metabolite profiles of red *Amaranthus* subjected to PAR (White) as control and monochromatic light treatments: deep blue, blue, green, amber, red, deep red, as well as far-red. Peaks corresponding to key metabolites-including organic acids like citric acid, malic acid, aconitic acid, amino acids like leucine, isoleucine, valine, phenylalanine, fatty acids like palmitic acid, stearic acid, phenolics like ferulic acid, caffeic acid and sugars like sucrose are annotated. Changes in peak intensities illustrate wavelength-dependent shifts in metabolic abundance. b) PCA score plots of red *Amaranthus* demonstrate clear and distinct clustering under different light wavelength with clear separation under far red and deep blue light wavelengths. c) Pathway enrichment analysis of metabolic profiles of the significant metabolic pathways using KEGG with an impact ≥ 0.1 for red *Amaranthus* seedlings under different light treatments. In a figure, each bubble depicts a metabolic pathway; the x-axis shows the pathway impact and y-axis shows the pathway enrichment. The key pathways impact and enrichment values are shown by the color and size, respectively.

### Intermediates of glyoxylate and dicarboxylate metabolism as well as TCA cycle showed wavelength and cultivar specific responses

In *Amaranthus* seedlings citric acid, aconitic acid, malic acid, oxalic acid, glycolic acid, glycine and glutamic acid which are the key metabolites of glyoxylate and dicarboxylate metabolism, were found to be affected under specific light wavelengths **(Figure 3, Figure 4)**. In green *Amaranthus*, malic acid levels decreased under far-red light treatment while all other light treatments resulted in non-significant levels **(Figure 3)**. The citric acid levels were significantly elevated under all light treatments except under far-red light treatment. The relative abundances of other metabolites of glyoxylate and dicarboxylate metabolism such as aconitic acid, oxalic acid, glycolic acid and glutamic acid were found to be decreased under both green and far-red light treatments. Oxalic acid and glutamic acid levels increased under amber light treatment, whereas glycine showed non-significant differences to all the light treatments. However, in red *Amaranthus*, the malic acid and citric acid levels decreased across all the light treatments **(Figure 4)**. Amber light elevated the levels of aconitic acid and oxalic acid significantly. The far-red light enhanced oxalic acid and glutamic acid levels. Glycolic acid showed non-significant variations among all the light treatments whereas glycine was found to be elevated under deep blue.

**Figure 3.**
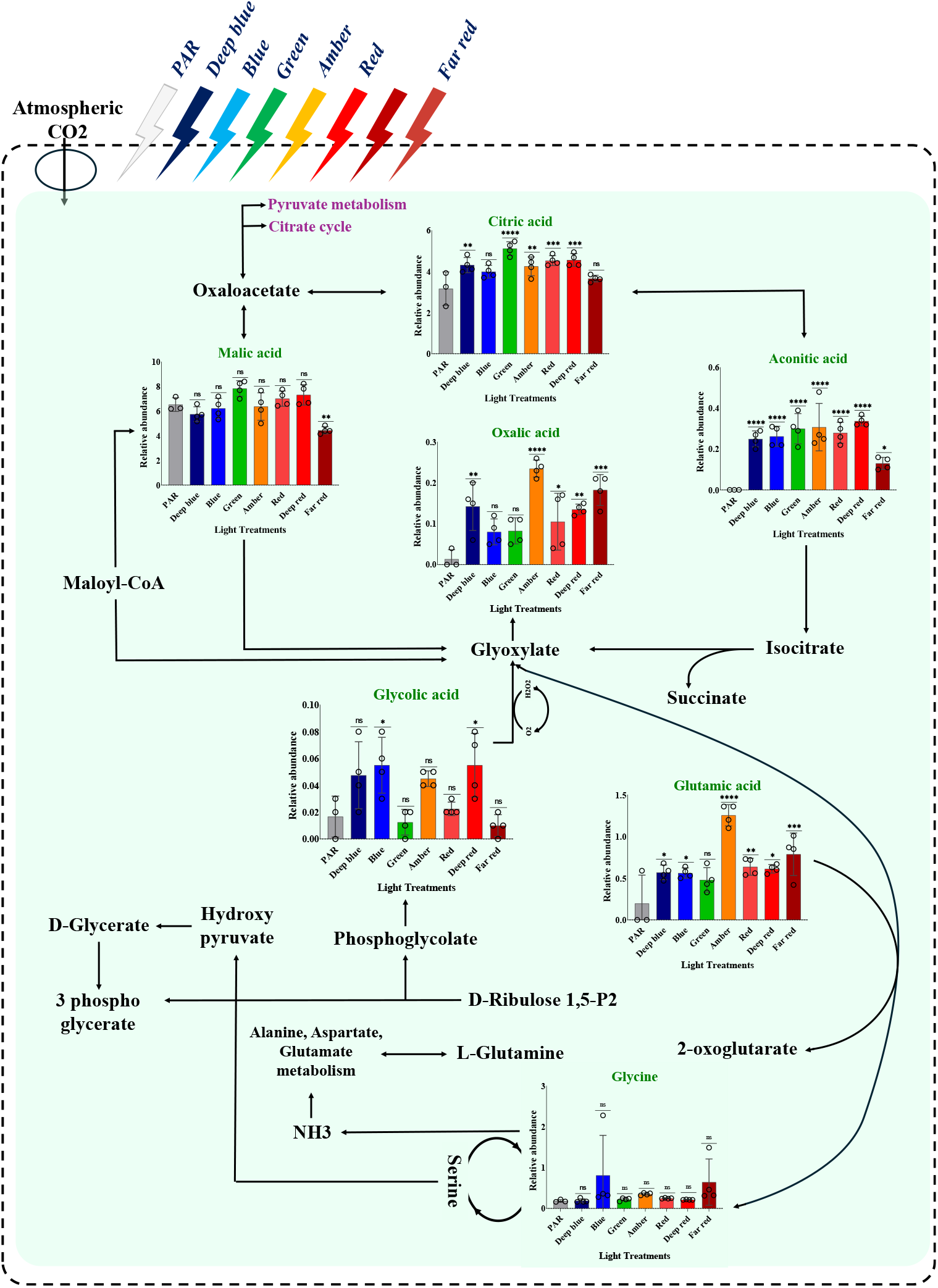
Pathway showing glyoxylate and dicarboxylate metabolism with relative abundances of metabolites under different light wavelength in green *Amaranthus*. The relative abundances of metabolites of glyoxylate and dicarboxylate metabolism plotted against white (PAR), deep blue, blue, green, amber, red, deep red, and far-red light wavelengths, are denoted by respective colour bars. Mean±SD represents the error bars with four replicates. One-way ANOVA and Dunnett’s multiple comparison test were used to determine statistical significance, with p value <0.05 = *, p value *<*0.01 = **, p value *<*0.001 = *** and p value <0.0001 = ****, considered significant and ns as non-significant. Bar graphs represent relative abundances of citric acid, aconitic acid, malic acid, oxalic acid, glycolic acid, glycine and glutamic acid under different light wavelengths in the green *Amaranthus*.

**Figure 4.**
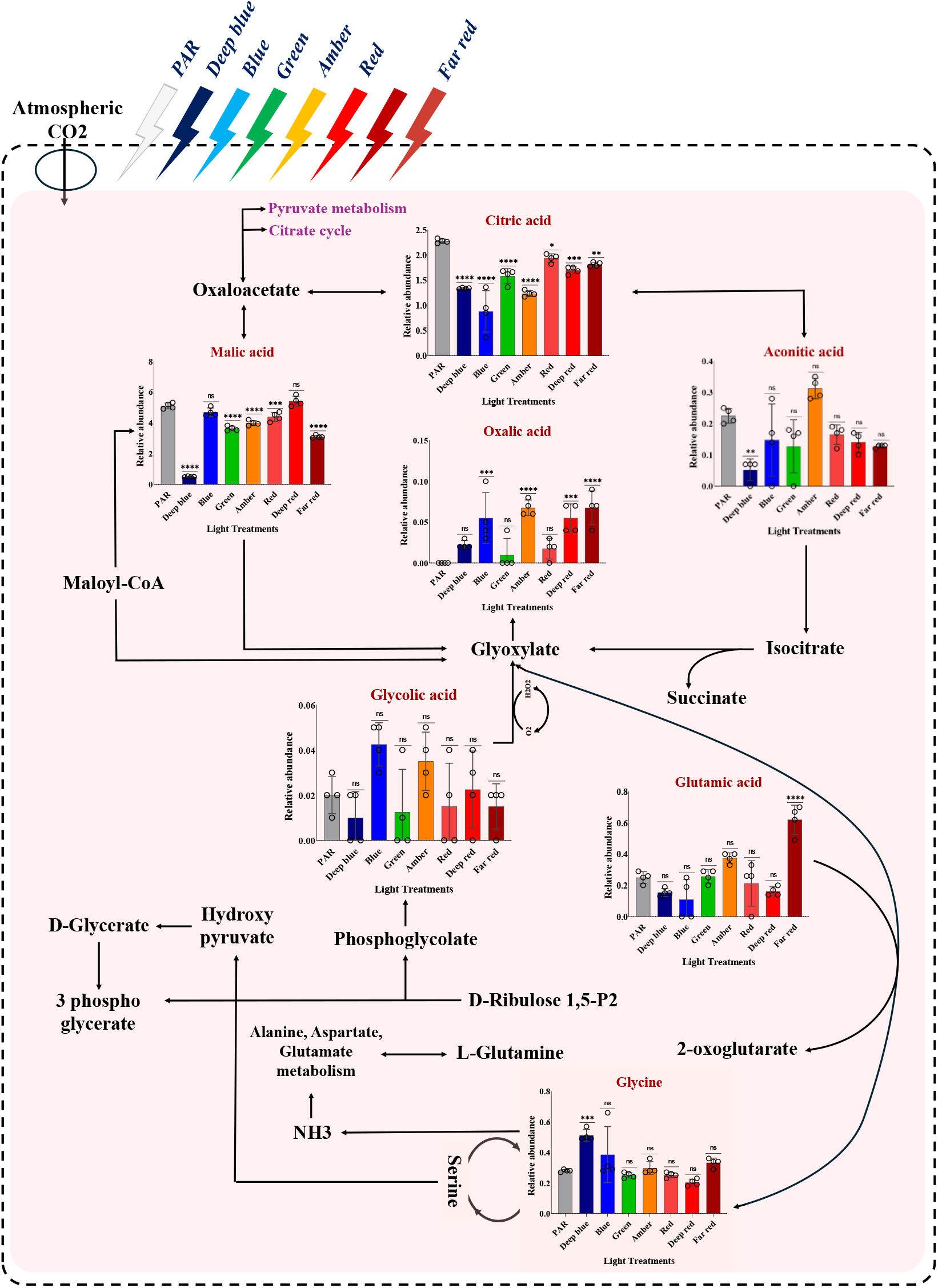
Pathway showing glyoxylate and dicarboxylate metabolism with relative abundances of metabolites under different light wavelength in red *Amaranthus*. The relative abundances of metabolites of glyoxylate and dicarboxylate metabolism plotted against white (PAR), deep blue, blue, green, amber, red, deep red, and far-red light wavelengths, are denoted by respective colour bars. One-way ANOVA and Dunnett’s multiple comparison test were used to determine statistical significance, with p value <0.05 = *, p value *<*0.01 = **, p value *<*0.001 = *** and p value <0.0001 = ****, considered significant and ns as non-significant. Mean±SD represents the error bars with four replicates. Bar graphs represent relative abundances of citric acid, aconitic acid, malic acid, oxalic acid, glycolic acid, glycine and glutamic acid under different light wavelengths in the red *Amaranthus*.

Glyoxylate and dicarboxylate metabolism is closely linked to TCA cycle and photorespiration. The metabolic analysis showed variable activities of TCA cycle and photorespiration under different wavelengths. Higher levels of the metabolites of TCA cycle were reported in plants grown under blue or white light compared to green and red light (Palma et al., 2022). Also in soybean cultivars, blue light increased the citrate and malate levels (Wang et al., 2025).Some of the metabolites of glyoxylate and dicarboxylate metabolism such as oxalic acid, citric acid, malic acid, glycine and glutamic acid are of health relevance. Oxalates are the metabolic end products which are considered as antinutrients which reduces the absorption of nutrients in humans. It’s excess consumption also leads to hyperoxaluria causing kidney stones (López-Moreno et al., 2022). In contrast citrate and malate whose levels are higher in *Amaranthus* seedlings can potentially inhibit the crystallization of oxalates decreasing the saturation of stone forming salts (Rodgers et al., 2014; Phillips et al., 2015). Further, glycine suppresses the calcium oxalate crystal accumulation in kidney by regulating urinary excretions of oxalate and citrate, while glutamate helps in nitrogen homeostasis (Lan et al., 2021; Loï and Cynober, 2022).

Both green and red *Amaranthus* seedlings on exposure to different light wavelengths showed distinct metabolic effects at the level of Glyoxylate and dicarboxylate metabolism thereby influencing the TCA cycle and photorespiratory pathways. This reflects wavelength and cultivar specific metabolic reprogramming of glyoxylate and dicarboxylate metabolism in *Amaranthus* seedling.

### Far-red light wavelength exposure enriched free amino acids of nutritional relevance in green and red *Amaranthus*

In plants, under certain conditions, metabolism can boost amino acid biosynthetic pathways which can in turn serve as precursors to many secondary metabolites with specialise functions (de Andrade et al., 2021). Previous research has shown the effect of blue, green, red as well as far-red wavelength treatment to amaranth microgreens leads to enhancement of phytochemicals (AY Ampim, 2020; Meas et al., 2020; Trandel-Hayse et al., 2025). In *Amaranthus*, most essential amino acids (EAAs) were enriched in seedlings grown under far-red light treatment in both the cultivars. EAAs cannot be synthesized by human body and are consumed as a supplement from the plant sources. EAAs such as branched chain amino acids (BCAAs), absorbed directly into muscle tissues and provides energy for physical activity whereas phenylalanine usually works on central nervous system and involved in the formation of neurotransmitters (Brestenský et al., 2015). There are many studies which have shown the beneficial impact of essential amino acid supplementation on human health and nutrition.

The increased levels of the phenylalanine (Phe) were observed in far red light grown seedlings compared to white light (**Figure 5a and 5e**). Isoleucine (Ile), leucine (Leu) and valine (Val) are crucial for plant growth, defense mechanisms, and energy supply (Gipson et al., 2017; Xing and Last, 2017; de Andrade et al., 2021). These were found to be strikingly higher in seedlings grown under far-red light in both cultivars (**Figure 5b, 5c, 5d, 5f, 5g, 5h**). Also, the lysine (Lys) as well as threonine (Thr) contents were significantly elevated by 35-fold and 11-fold respectively in green *Amaranthus* seedlings under far-red wavelength **(Supplementary figure 2)**. Followed by far-red, in amber light also lysine, threonine and branched chain amino acids were found to be elevated. These amino acids are known to be present in very high concentration in both green and red leafy cultivars than in grain type cultivars.

**Figure 5.**
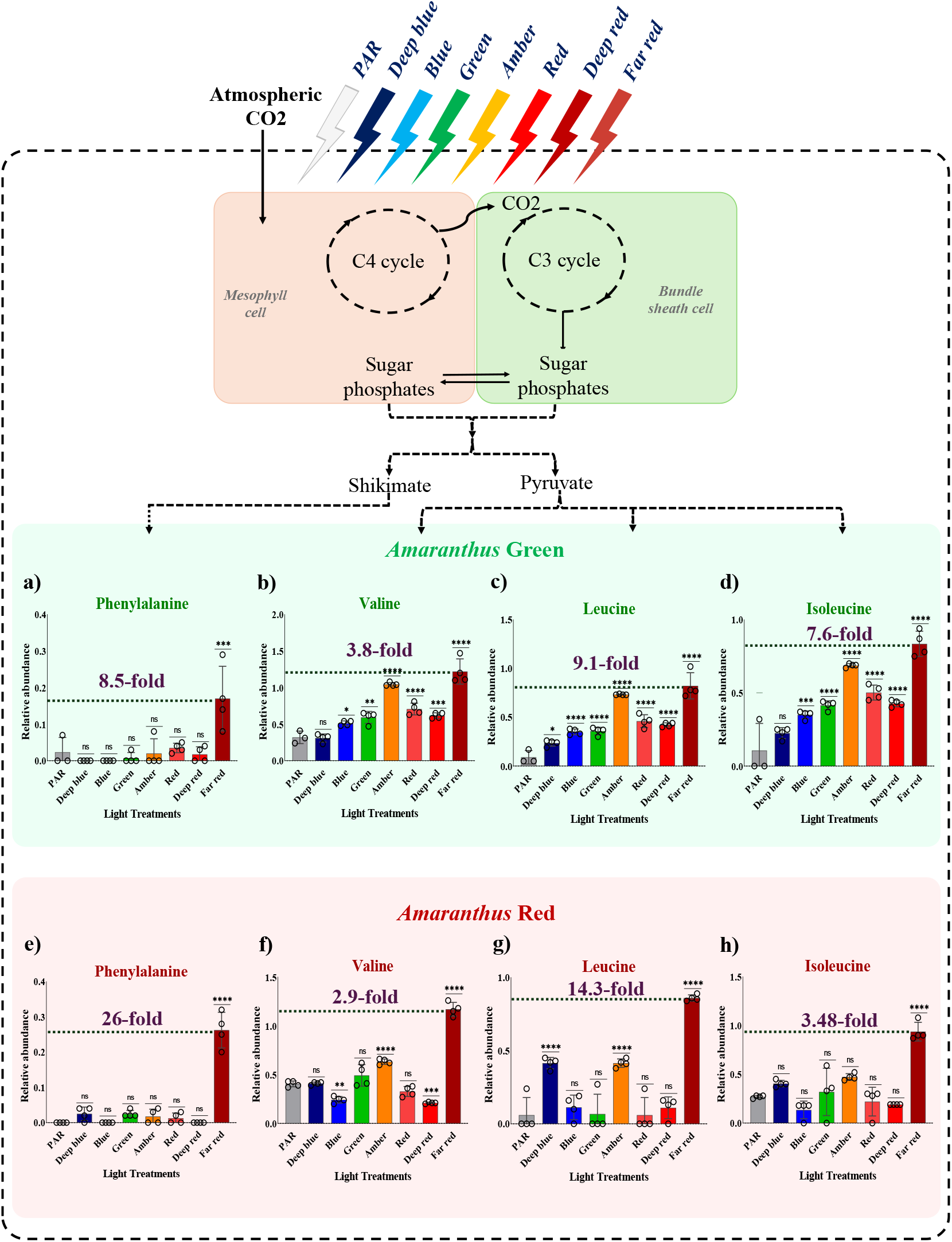
Relative abundances of essential amino acids show targeted enrichment under far-red light wavelengths in green and red *Amaranthus* cultivars. The relative abundances of essential amino acids plotted against white (PAR), deep blue, blue, green, amber, red, deep red, and far-red light wavelengths, are denoted by respective colour bars. One-way ANOVA and Dunnett’s multiple comparison test were used to determine statistical significance, with p value <0.05 = *, p value *<*0.01 = **, p value *<*0.001 = *** and p value <0.0001 = ****, considered significant and ns as non-significant. Mean±SD represents the error bars with four replicates. a, b, c and d represent the relative abundances of phenylalanine, valine, leucine, isoleucine under different light wavelengths in the green *Amaranthus*, whereas, e, f, g and h represent the relative abundances of phenylalanine, valine, leucine, isoleucine under different light wavelengths in red *Amaranthus*.

Studies across various species, including wheat, barley, *Arabidopsis, Auricularia cornea* have shown that LED treatments can modulate amino acid content (Koga et al., 2013; Yang et al., 2016; Toldi et al., 2019; Ye et al., 2025). In wheat, levels of the specific amino acids were variably influenced whereas, barley demonstrated a consistent increase, highlighting the unique impact of monochromatic red wavelength. Also, in plants like chinese kale and leafy vegetables, supplementation with far-red light enhanced the accumulation of free amino acids, particularly BCAAs along with phenylalanine (Li et al., 2021; Frutos-Totosa et al., 2023). Studies on various plants have reported an increased levels of phenylalanine under far-red light which negatively impacts flavonoid accumulation, the end products of the phenylpropanoid pathway originating from phenylalanine (Liu et al., 2015; Yeow et al., 2020). Far-red light affects metabolism via phytochrome signalling. Red/far-red light ratio influences phytochrome photo equilibrium, which controls physiological and developmental responses like metabolic allocation and enzyme activity (Li et al., 2021). This signalling results in broad physiological as well as metabolic changes, including modulating amino acid biosynthesis and allocation. The observed increase in phenylalanine levels compared to phenolic acids under far red light might be an adaptive response in *Amaranthus* cultivars. Furthermore, the shift in metabolism away from secondary metabolites toward amino acid accumulation under far-red light suggests a reallocation of carbon sources, potentially favouring the synthesis and storage of BCAAs and other amino acids.

### Phenolics showed cultivar specific responses to different light wavelengths in *Amaranthus tricolour*

The phenolic compounds, caffeic acid and ferulic acid showed major variations in the green and red *Amaranthus* seedlings under different light wavelengths. Phenolic acids are the secondary metabolites which can be strongly influenced by light quality and intensity, with different wavelengths modulating their biosynthesis (Ouzounis et al., 2014; Bantis et al., 2018; Xiong et al., 2025). They can function as antioxidants, regulates physiological processes and modulates stress responsive pathways in plants (Poggioni et al., 2022; Saini et al., 2024; Q. Sun et al., 2024). Phenolics are also reported for their role in human health as antioxidant, anti-inflammatory and disease preventive properties (Cao et al., 2023; El-Wekil et al., 2025).

Caffeic acid was elevated under the exposure of green light in the green *Amaranthus* and was detected to be about 2.73-fold higher compared to the white light exposure (**Figure 6a**). Compared to other wavelengths, green light is inefficient in enhancing chlorophyll content in plants. Chlorophyll mostly absorbs wavelength in the red (∼660 nm) as well as blue (∼450 nm) spectra, whereas green light is absorbed to a lesser extent, predominantly being reflected, which gives the characteristic colour to plants. However, research indicates in spite of fact that, green light is less effective at photosynthesis and indirectly facilitates chlorophyll production, it penetrates deeper into leaves and cellular layers (Leite et al., 2025). In certain instances, this effect can potentially enhance total photosynthetic efficiency by balancing red and blue light. Plantlets of *Aeollanthus suaveolens* cultivated in green wavelength exhibited significant levels of carotenoid and chlorophyll a, b (Araújo et al., 2021). The green wavelength can penetrate plant canopy more profoundly than blue and red wavelength, influencing growth along with the production of bioactive chemicals through both cryptochrome-dependent and independent mechanisms (J. Sun et al., 1998; Dou et al., 2019). The lowest accumulation of phenolic compounds was observed in sow thistle plantlets grown under monochromatic green light (Leite et al., 2025).

**Figure 6.**
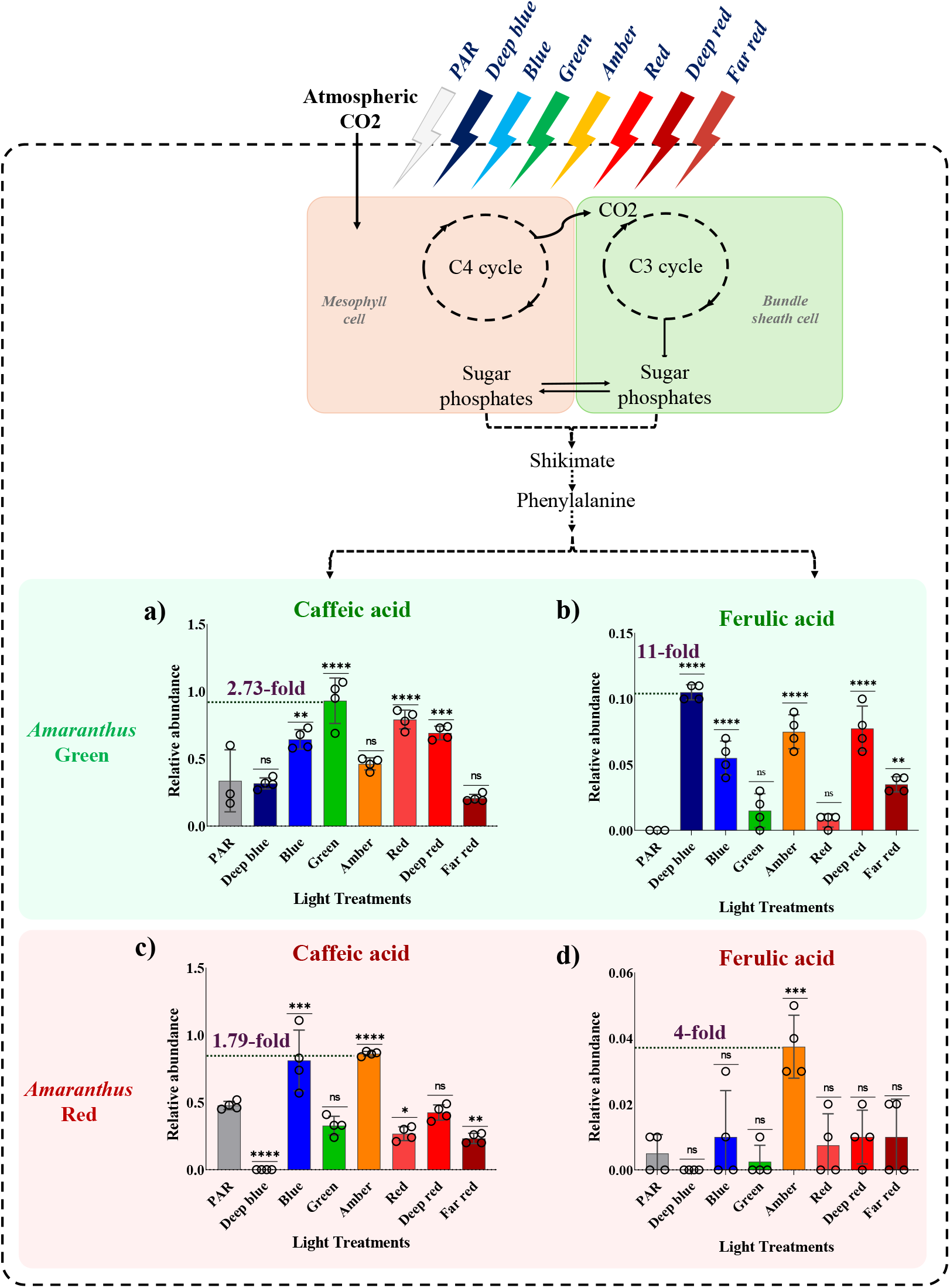
Relative abundances of phenolic acids showing targeted enrichment under variable light wavelength in green and red *Amaranthus* cultivars. The relative abundances of phenolic acids plotted against white (PAR), deep blue, blue, green, amber, red, deep red, and far-red light wavelengths, are denoted by respective colour bars. One-way ANOVA and Dunnett’s multiple comparison test were used to determine statistical significance, with p value <0.05 = *, p value *<*0.01 = **, p value *<*0.001 = *** and p value <0.0001 = ****, considered significant and ns as non-significant. Mean±SD represents the error bars with four replicates. a and b represent the relative abundances of caffeic and ferulic acid under different light wavelengths in the green *Amaranthus*, whereas c and d represent the relative abundances of caffeic and ferulic acid under different light wavelengths in red *Amaranthus*.

In green *Amaranthus* the ferulic acid was found to be 11-fold elevated under deep blue light treatment (**Figure 6b**). Deep blue light due to its high energy and short wavelength can induce the synthesis of phenolic compounds in plants. However, it is crucial to understand that the effects might be species or cultivars-dependent, with different species exhibiting varying responses in phytochemical accumulation. We observed that red *Amaranthus* on exposure to deep blue has lowered the levels of ferulic acid **(Figure 6d)**. However, it is of interest to observe that in red *Amaranthus*, both caffeic and ferulic acid was found to be elevated under amber light treatment. Caffeic acid was detected 1.79-fold higher and ferulic acid was 4-fold higher under amber light treatment when compared to white light (**Figure 6c and 6d**).

## Conclusion

Nutrient-dense food sources which can be rapidly and locally cultivated are essential for the overall health of consumers. Edible microgreens are considered nutritionally dense, possess a brief growth cycle, and necessitate minimal resources, which makes them excellent options for promoting human health (Dereje et al., 2023; Poudel et al., 2024; Trandel-Hayse et al., 2025). The present study investigated how different monochromatic light wavelengths of visible range affects the metabolic profiles of green and red leafy cultivars of *Amaranthus tricolor*. GC-MS based metabolite profiling of *Amaranthus* cultivars exposed to deep blue, blue, green, amber, red, deep red, far red, and white light as a control (PAR region; 400-700 nm) revealed wavelength induced metabolic responses. Under far red-light wavelengths enhanced levels of the branched chain amino acids and phenylalanine is observed in both the cultivars. However, variable metabolic response in both the cultivars was observed in relation the accumulation of phenolic acids. In the green cultivars, caffeic acid and ferulic acid levels were elevated under the green light and deep blue light respectively. In contrast, both caffeic and ferulic acids were elevated under amber light in the red cultivars. This reprogramming suggests the metabolic adjustments of *Amaranthus* seedlings under different light wavelengths.

Our findings highlight that light treatment with specific monochromatic wavelengths reprograms the metabolism in *Amaranthus* seedlings and provides cues towards targeted enrichment of metabolites. Far-red light can induce targeted enrichment of essential amino acids for both cultivars. In contrast, phenolic acids showed wavelength and cultivars specific responses. Green and deep blue light showed enrichment of caffeic and ferulic acid respectively in green *Amaranthus* seedlings, whereas amber light showed enrichment of both phenolic acids in red *Amaranthus* seedlings. Overall, this study highlights the potential of wavelength specific light modulation for targeted enrichment of metabolites in green and red *Amaranthus* seedlings, offering strategies for improvement of its nutritional value in controlled environmental conditions.

## Supplementary data

Additional information included here

**Supplementary Figure 1**: Experimental design of green and red *Amaranthus* metabolomics under different light wavelengths

**Supplementary Figure 2**: Relative abundances of lysine and threonine showing targeted enrichment under far-red light wavelength in green *Amaranthus*

## Acknowledgements

SKM acknowledges the funding agency Science and Engineering Research Board (SERB) for Early career research funding (File No: ECR/2016/001176). SSP acknowledges Department of Science and Technology (DST) for her Inspire PhD fellowship. NJ acknowledge L’institut Agro Montpellier, France for his Masters. ML and YP acknowledge Ministry of Education for their PhD fellowships. All authors acknowledge IIT Mandi for the facilities and support.

## Author contribution

SKM and ML conceptualised the study. NJ, YP and ML performed the experiments. SSP and SKM analysed the data. SSP made all the figures and prepared original draft. Also, all authors reviewed as well as edited the manuscript. SKM did the research supervision and also acquired the funding.

## Data Availability

The manuscript figures and supplementary data contain all supporting information. It will be provided on request.

