## Supplementary figures for "Wavelength induced cultivar specific enrichment of essential amino acids and phenolics in *Amaranthus tricolor*"

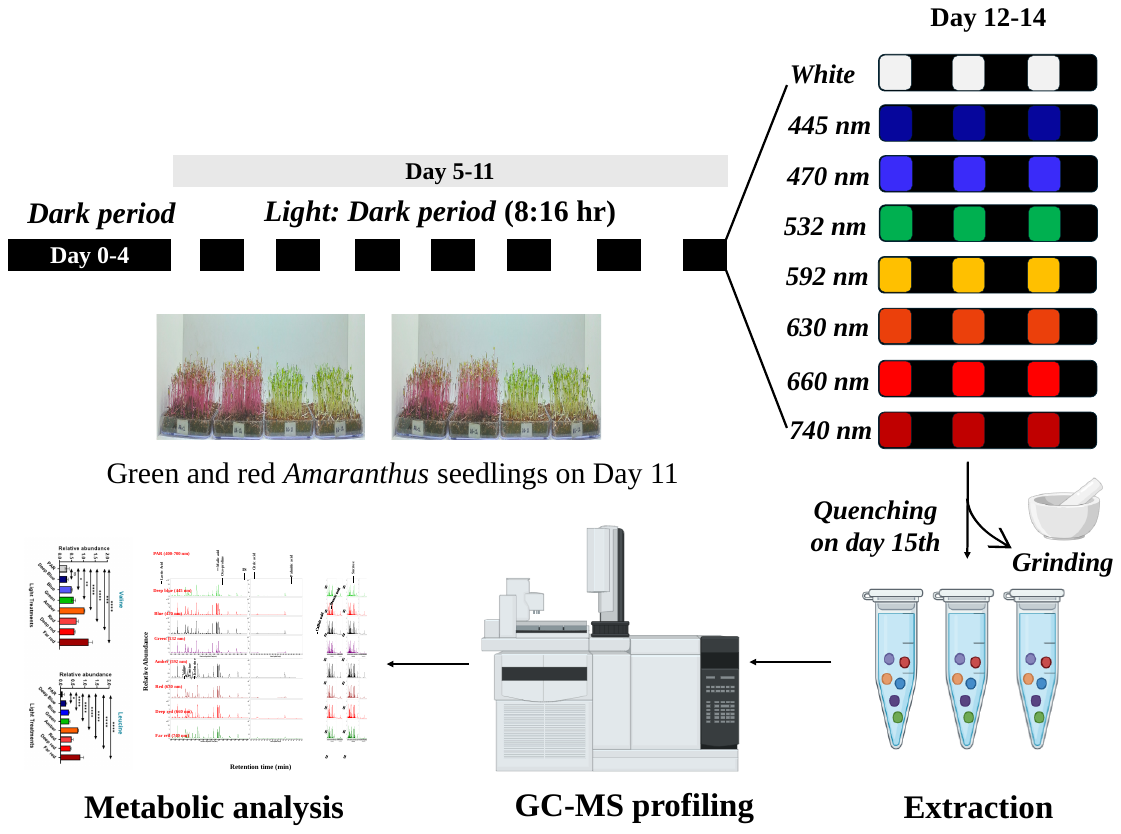

**Supplementary figure 1. Experimental design of green and red *Amaranthus* metabolomics under different light wavelengths.** Seedlings were first maintained in darkness (day 0-4) and then shifted under an 8:16 hr light-dark photoperiod (D5-D11). On days D12-D14, plants were exposed to different light wavelengths with PAR (White) light as control and seven monochromatic wavelengths (445, 470, 532, 592, 630, 660, and 740 nm). On Day 15, the tissues were rapidly quenched with liquid nitrogen, ground, and subjected to metabolite extraction. Extracts were analyzed using GC-MS, and the resulting chromatographic and quantitative data were used for metabolic analysis across light treatments and cultivars.

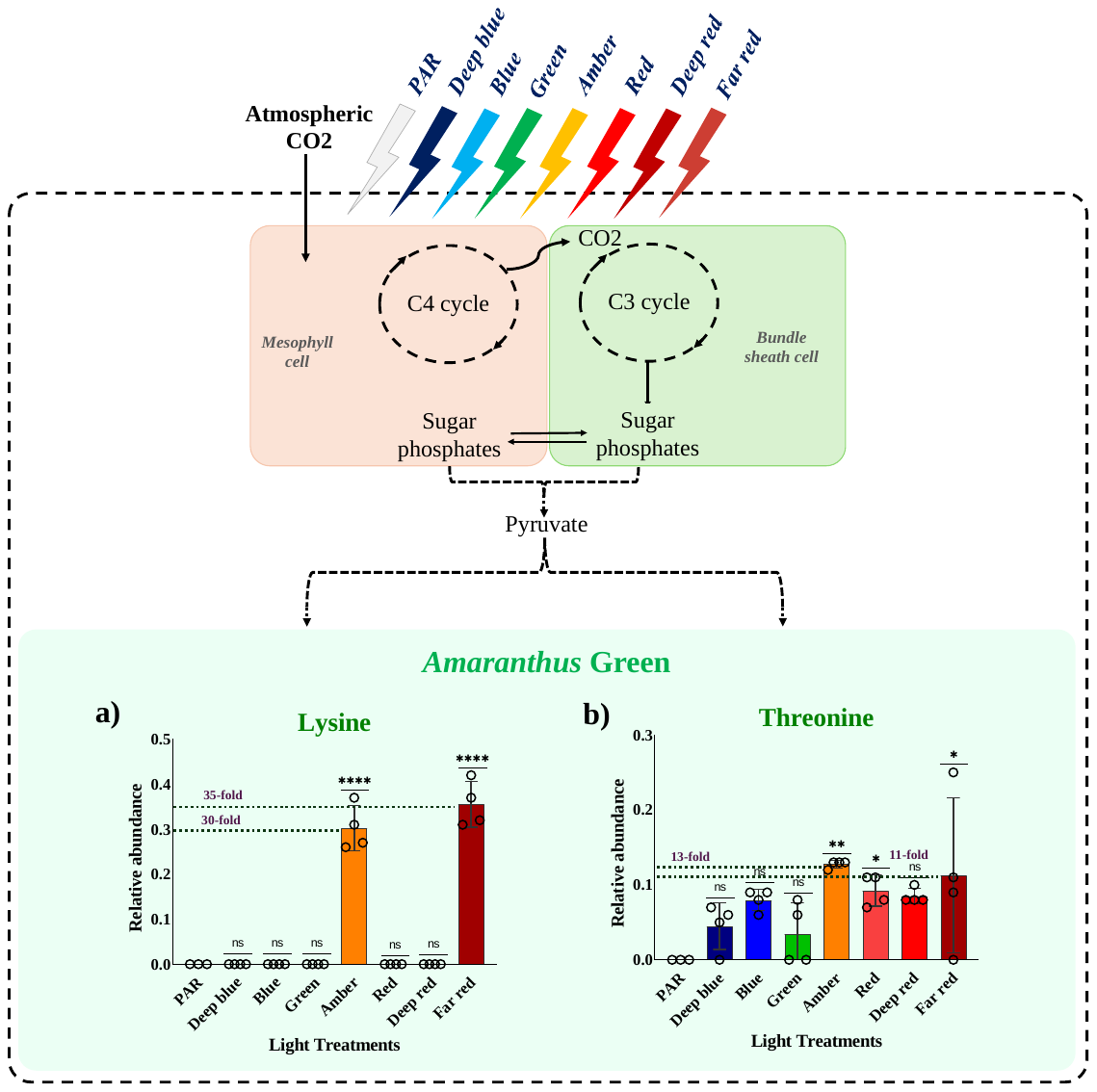

**Supplementary figure 2. Relative abundances of lysine and threonine showing targeted enrichment under far-red light wavelength in green *Amaranthus.*** The relative abundances of essential amino acids plotted against white (PAR), deep blue, blue, green, amber, red, deep red, and far-red light wavelengths, are denoted by respective colour bars. One-way ANOVA and Dunnett’s multiple comparison test were used to determine statistical significance, with p value <0.05 = *, p value *<*0.01 = **, p value *<*0.001 = *** and p value <0.0001 = ****, considered significant and ns as non-significant. Mean±SD represents the error bars with four replicates. a represent the relative abundances of lysine whereas, b represents relative abundances of threonine under different light wavelengths in green *Amaranthus*.
